# The Two Frontiers Project Field Handbook and OpenTools: Standardizing microbial fieldwork for biobank-scale sequencing and culturomics

**DOI:** 10.64898/2025.12.25.695958

**Authors:** Krista A Ryon, James R Henriksen, April Johns, Sara C Diana, Victor Boddy, Gaby E. Carpenter, Erin Miller, Ben Kent, Raquel Peixoto, Braden T Tierney

## Abstract

From medicines to materials, our planet’s microbial diversity comprises an enormous wellspring of biotechnological potential. For centuries, microbiologists have developed tools for interrogating microbial function, ranging from microscopy and culturing to, more recently, metagenomics. However, deploying these tools during fieldwork requires substantial forward planning, interdisciplinary technical expertise, and plans for navigating permitting and the ethical implications of bioprospecting. To address these challenges, we built The Two Frontiers Project Handbook and OpenTools Resource, which aggregates our expertise in high-throughput sampling, sequencing, and culturing of microbes from thousands of samples. We provide our full suite of fieldwork methods as well as relevant software and hardware. We lay our standards for team roles and construction, general expedition planning, sample transport, permitting, and numerous other key aspects of executing a successful field campaign. The version-controlled resource is available at https://two-frontiers-project.github.io/ and is open for non-commercial use.

## Introduction

Earth’s biomes contain an estimated 5x10^28^ microbial cells that together comprise millions of species and billions of genes^1,2^. Yet, most microbes are still considered as “dark matter”, as only a small fraction of this immense diversity has been characterized through DNA sequencing and other approaches^3–5^. Even this limited exploration has yielded transformative biotechnological breakthroughs, including CRISPR, Polymerase Chain Reaction, antibiotics, other novel drugs, carbon sequestering organisms, and probiotics for coral reef health^6–10^. There is no question that the unsequenced and uncultured majority contain manifold, highly impactful discoveries just waiting to be found. Indeed, the recent launch of the microbial conservation specialist group by the International Union for Conservation of Nature (IUCN) also highlights a growing awareness of this diversity, as well as the need to protect, catalog, and conserve it in biobanks^11^.

However, accessing this diversity at scale remains technically challenging and requires a suite of multidisciplinary, complementary approaches. These fall into two general categories: (1) the “traditional” tools of microbiology (e.g., microscopy, culturing, numerical taxonomy) and (2) high-throughput molecular approaches (e.g., shotgun metagenomics). Any single method deployed alone provides only a partial view. Metagenomics, for example, provides a comprehensive snapshot of the microbial community, revealing its taxonomic composition and genomic potential (i.e., the organisms and genes present), including even those that have yet to be cultured. However, without the tools of microbiology and genetic engineering to isolate, culture, and characterize microbial function, such insights remain largely theoretical and difficult to translate into practical applications, like bioproduction or ecosystem restoration^8,9^. Culture-based microbiology directly addresses this gap, although when applied in isolation, it often lacks direction, as it provides little guidance on which organisms or functions to pursue. In a sense, next-generation sequencing provides a map, while traditional microbiology supplies the means to explore it, and these approaches can be further enhanced by combining culturing and metagenomic insights (i.e., through culturomics) with in-situ cultivation approaches^12,13^.

Additionally, numerous ethical considerations must be addressed when querying global microbial diversity to discover useful functions. Bioprospecting (i.e., the practice of sampling extreme or diverse environments around the world in pursuit of valuable microbial technologies) has a complex and troubling history^14,15^. While previous expeditions have led to transformative discoveries, they have often done so without fair recognition or compensation for local or Indigenous communities. The benefits have historically flowed almost exclusively to the collecting parties – typically researchers from wealthy, historically colonial nations^16^.

To address these and other inequities, numerous regulations and agreements have been established to promote both ethical and equitable bioprospecting. The Nagoya Protocol (NP) on Access and Benefit-Sharing (ABS), adopted under the Convention on Biological Diversity, provides a legal framework ensuring that benefits arising from the utilization of genetic resources are shared fairly with the countries and communities from which those resources originate^17,18^. Similarly, the recently ratified High Seas Treaty (formally the Agreement on Biodiversity Beyond National Jurisdiction, or BBNJ Agreement) extends these principles to marine genetic resources in international waters, establishing a mechanism for benefit sharing, capacity building, and scientific collaboration in regions beyond any national jurisdiction^19^.

Despite these frameworks, researchers still face a complex and often nebulous network of international, governmental, and regional permitting systems that are difficult to navigate efficiently^20^. Moreover, these formal agreements and treaties do not always translate into clear, locally implementable benefit-sharing practices or reflect the priorities of all affected stakeholders^21^. Therefore, there is a pressing need for practical, accessible frameworks and resource hubs to guide field scientists in conducting both scientifically rigorous and ethically responsible bioprospecting expeditions.

As a result, there is still a need for streamlined and publicly available frameworks for harmonizing modern and traditional microbiological approaches for ethical bioprospecting. Standards, such as those from the National Microbiome Data Collective, provide guidance for data storage, metadata collection, and analysis^22,23^. However, universal fieldwork protocols that integrate these practices across methodologies are currently not yet consolidated into a single, openly available resource.

To address this gap, The Two Frontiers Project (2FP) has spent the past three years developing standardized, low-cost workflows for exploring, understanding, and leveraging microbial diversity. Specifically, we have endeavoured to build workflows that enable the integration of traditional microbiology fieldwork tools with modern methods of next-generation sequencing, all while accounting for the ethical and regulatory complexities of international sampling. Here, we present for the scientific community the assemblage of these methods and resources as The 2FP Field Handbook and OpenTools Resource.

## Results

### The Two Frontiers Handbook: Overview, design principles, and usage

The goal of The Two Frontiers Handbook and OpenTools Resource (i.e., the 2FP Handbook) is to unify and simplify best practices for highly scalable (e.g., to thousands of samples) (1) DNA sequencing, (2) cryopreservation, and (3) culturing diverse microbial communities. These three experimental practices have divergent and, at times, conflicting requirements; as a result, they are rarely executed simultaneously at large scale. For example, buffers for DNA preservation often disrupt cell viability, preventing downstream culturing assays. Conversely, cryopreservation mandates maintaining a cold chain, ideally with dry ice. Additionally, viable cells versus DNA require distinct import/export and ethical considerations and processing methods in-field.

We developed the 2FP Handbook to provide a point of reference for harmonizing the, at times discordant requirements of field microbiology. It is available at https://two-frontiers-project.github.io/#/ for research-use only under a non-commercial license. The Handbook itself is broken into 10 sections, each of which comprises a sequential step in the expedition planning process (Table 1). We developed this knowledge base and protocol over the past three years of sampling, sequencing, culturing, and biobanking thousands of samples from around the globe from both terrestrial and aquatic ecosystems (Figure 1).

**Table 1.**
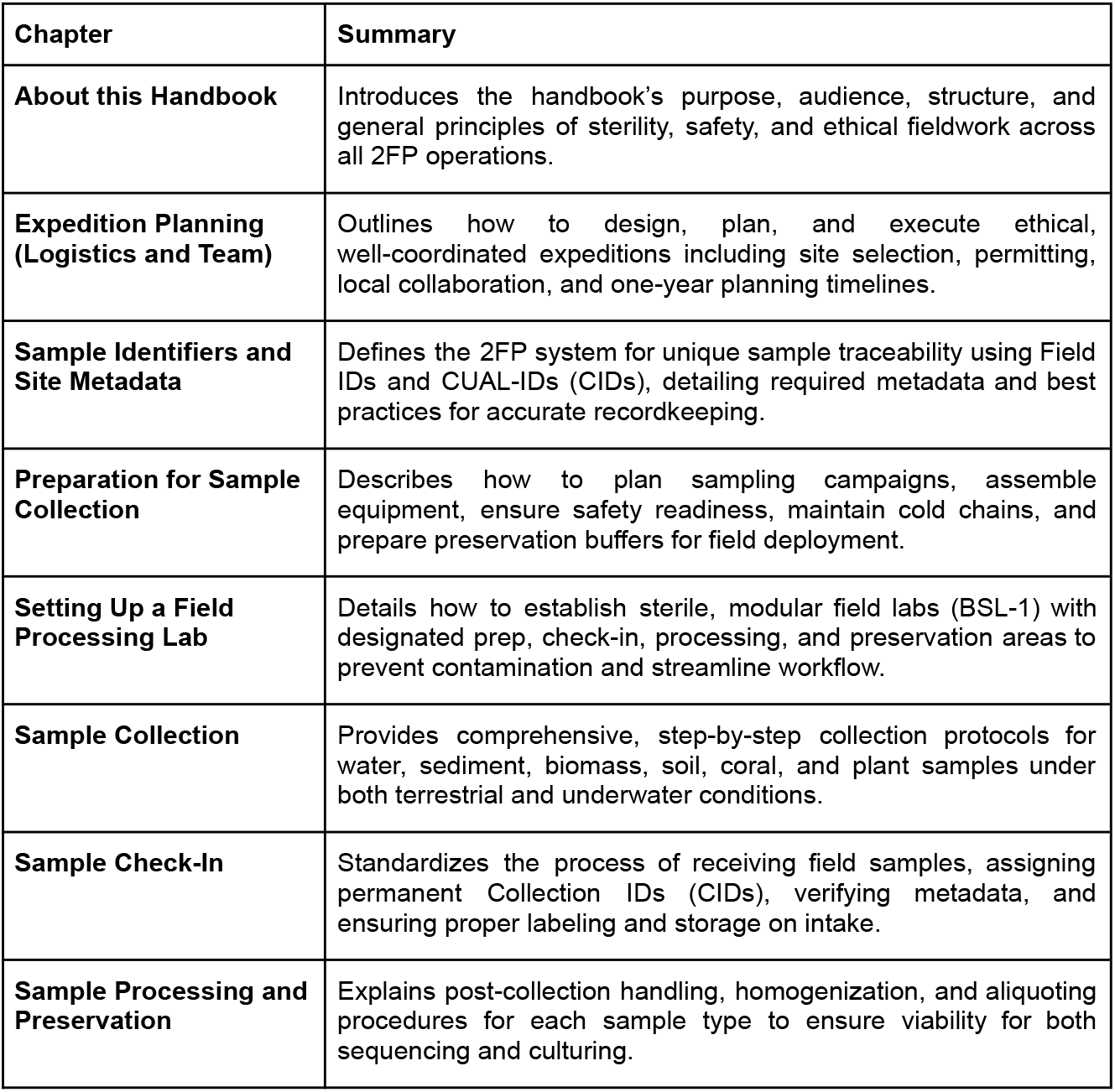
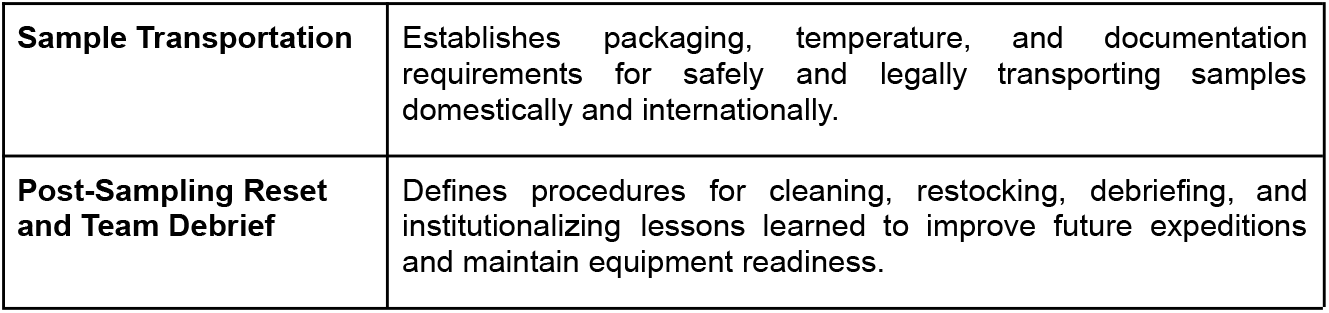
Handbook chapters.

| Chapter | Summary |
| --- | --- |
| **About this Handbook** | Introduces the handbook's purpose, audience, structure, and general principles of sterility, safety, and ethical fieldwork across all 2FP operations. |
| **Expedition Planning (Logistics and Team)** | Outlines how to design, plan, and execute ethical, well-coordinated expeditions including site selection, permitting, local collaboration, and one-year planning timelines. |
| **Sample Identifiers and Site Metadata** | Defines the 2FP system for unique sample traceability using Field IDs and CUAL-IDs (CIDs), detailing required metadata and best practices for accurate recordkeeping. |
| **Preparation for Sample Collection** | Describes how to plan sampling campaigns, assemble equipment, ensure safety readiness, maintain cold chains, and prepare preservation buffers for field deployment. |
| **Setting Up a Field Processing Lab** | Details how to establish sterile, modular field labs (BSL-1) with designated prep, check-in, processing, and preservation areas to prevent contamination and streamline workflow. |
| **Sample Collection** | Provides comprehensive, step-by-step collection protocols for water, sediment, biomass, soil, coral, and plant samples under both terrestrial and underwater conditions. |
| **Sample Check-In** | Standardizes the process of receiving field samples, assigning permanent Collection IDs (CIDs), verifying metadata, and ensuring proper labeling and storage on intake. |
| **Sample Processing and Preservation** | Explains post-collection handling, homogenization, and aliquoting procedures for each sample type to ensure viability for both sequencing and culturing. |
| **Sample Transportation** | Establishes packaging, temperature, and documentation requirements for safely and legally transporting samples domestically and internationally. |
| **Post-Sampling Reset and Team Debrief** | Defines procedures for cleaning, restocking, debriefing, and institutionalizing lessons learned to improve future expeditions and maintain equipment readiness. |

**Figure 1.**
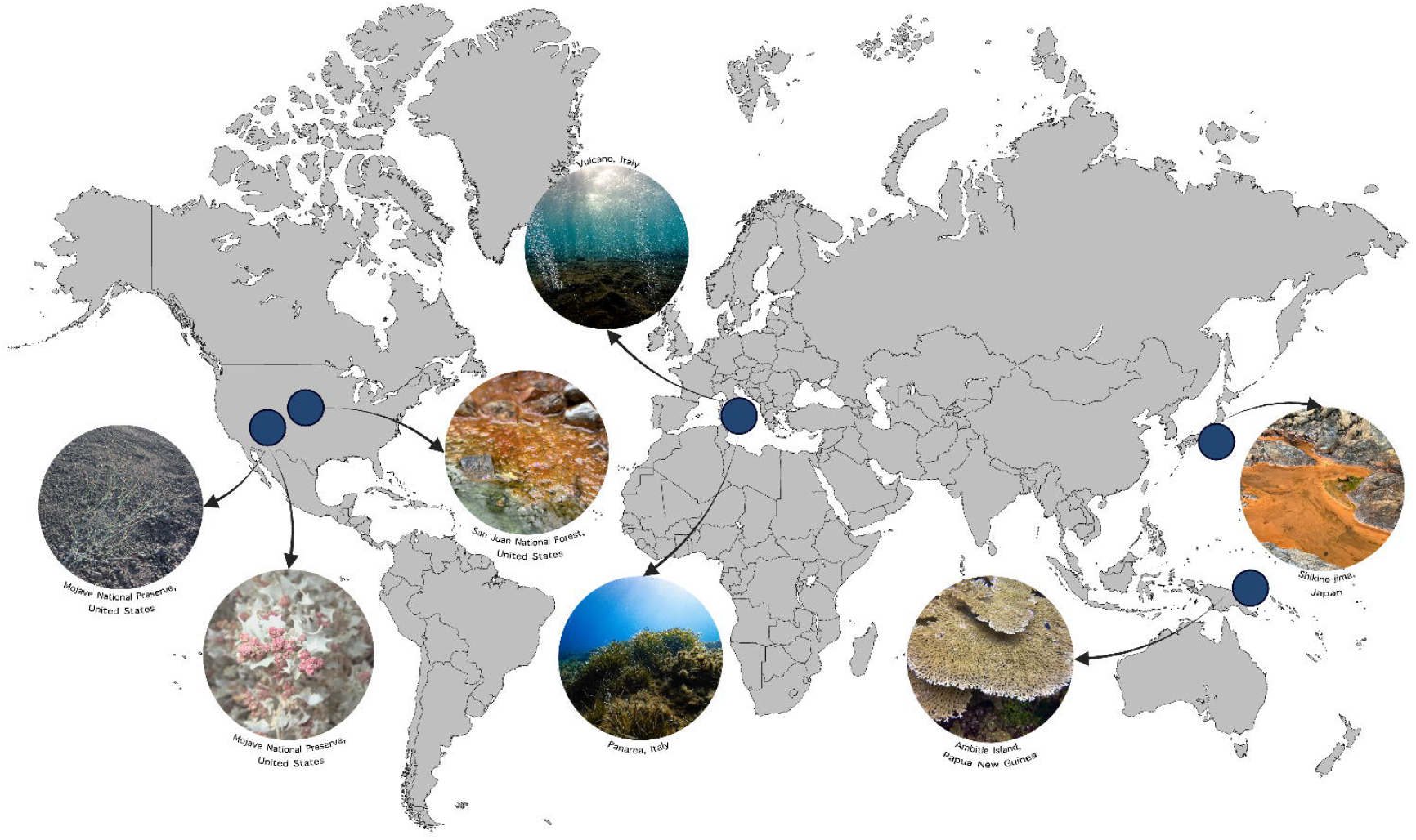
Examples of areas sampled and sample types collected by The Two Frontiers Project.

The structure of this Handbook draws inspiration from the instructional design principles used in the *Instructor Manual* of the Professional Association of Diving Instructors (PADI), emphasizing clarity, modularity, and practicality as a reference. While distinct in purpose and content, it similarly serves as a systematic guide for trained users to consult as questions arise in the field. Adapting the concept of waterproof “slates” used in diving instruction, we developed a series of concise, durable field protocols (Figure 2–3) that can be printed on weather-resistant material and used directly during sampling. Current versions include slates for metadata collection, terrestrial and aquatic sampling, processing, and transport. All are version-controlled and publicly available through our GitHub repository on the 2FP OpenTools site (https://github.com/two-frontiers-project/2FP-sampling-slates).

**Figure 2.**
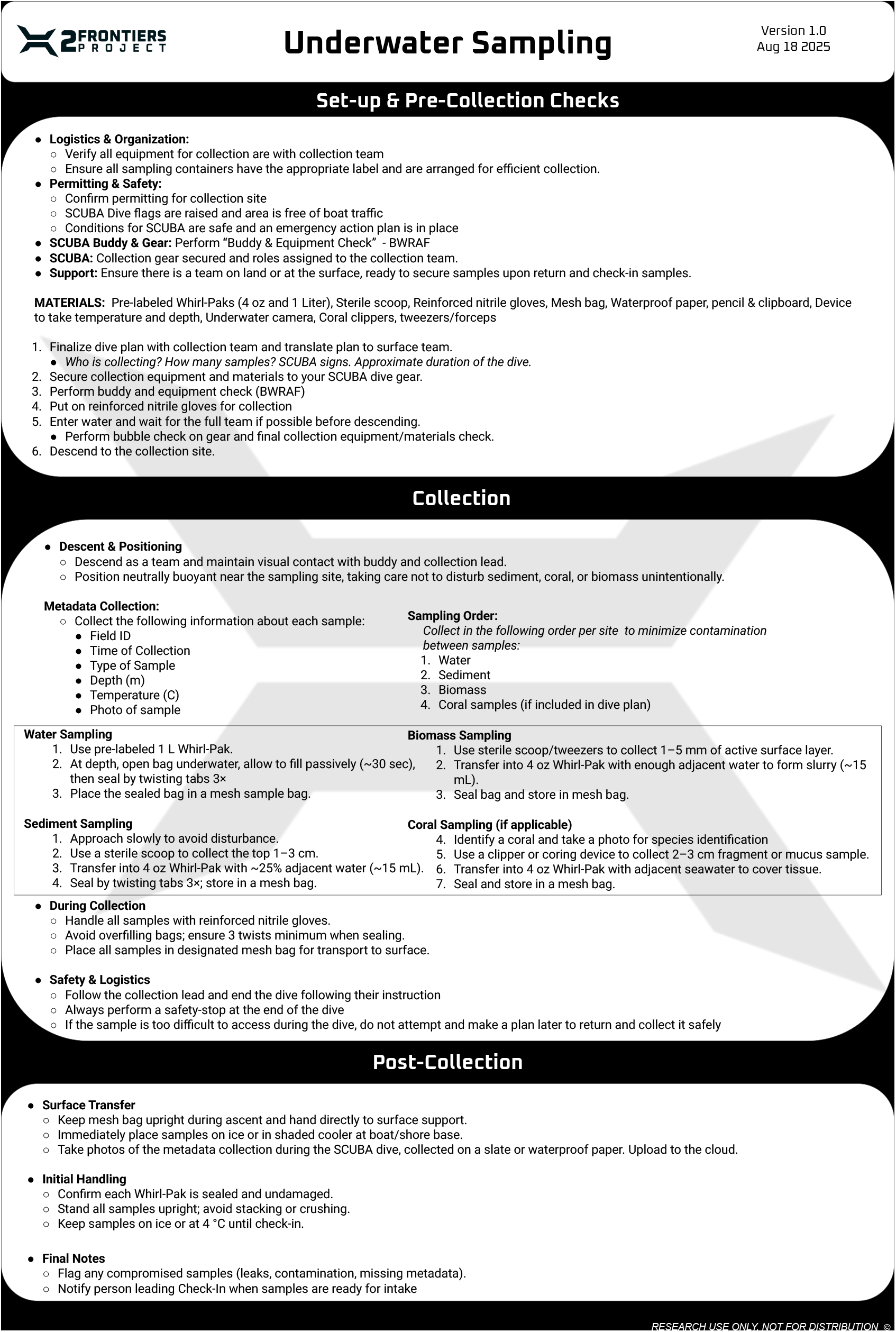
The 2FP underwater sampling slate.

**Figure 3.**
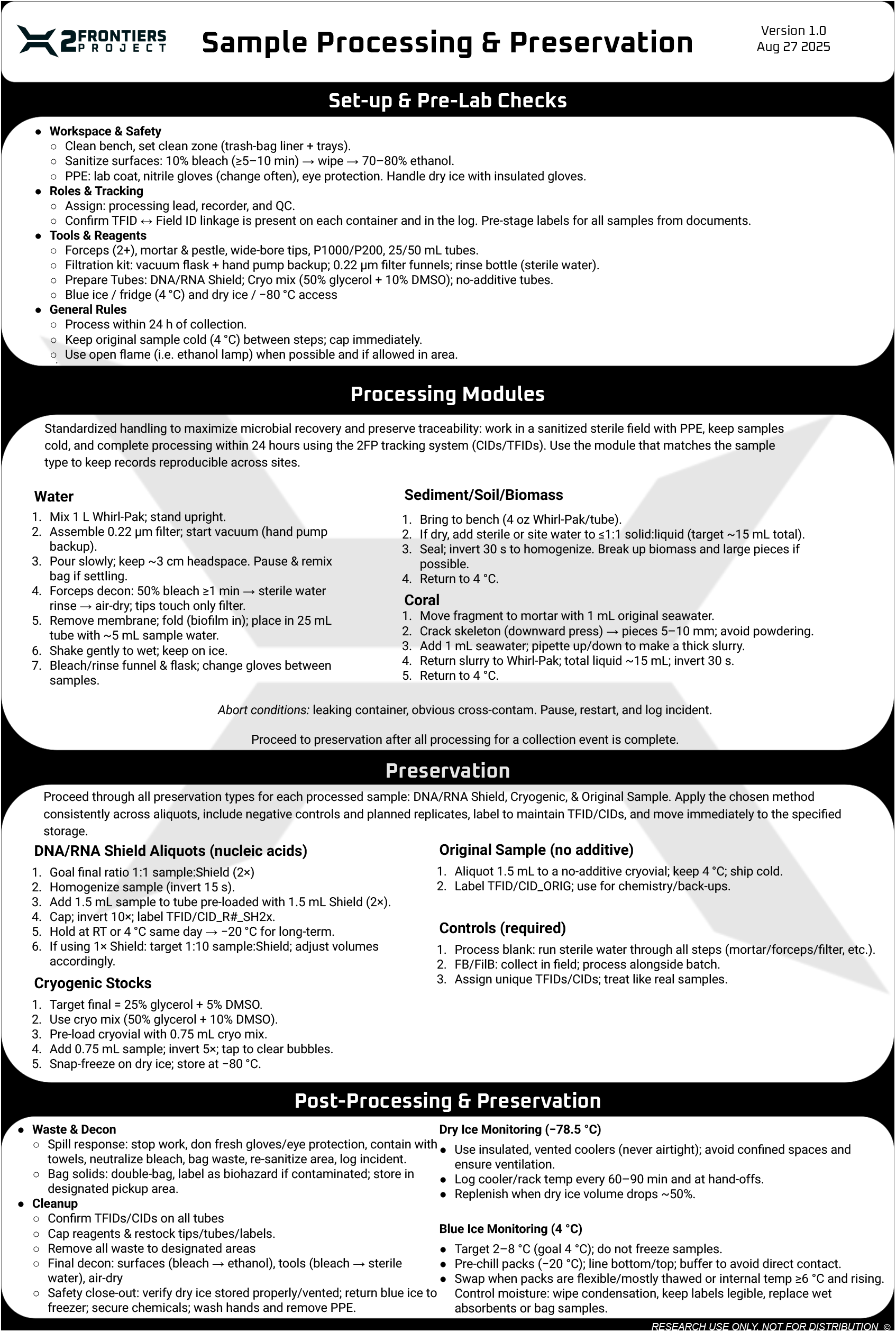
The 2FP sampling processing slate.

### Pre-sampling planning: permitting, ethics, logistics, and team roles

The first Handbook sections cover forward planning for sampling outings. These include equipment acquisition and protocol design, assembling a team with the appropriate expertise and dispositions, and considering the ethical and permitting implications of sample collection their downstream use. We outline a one-year planning cycle that we try to implement for the majority of large-scale or international efforts. For smaller, one-off collections (e.g., single-day trips) that do not involve import/export restricted samples or complex protocols (e.g., field sequencing or on-site culturing media design), timelines can be compressed to a matter of months. We specify strategies for building an effective field team with an appropriate distribution of skillsets and roles. Finally, this section also covers safety guidelines and provides protocols for preparing various preservatives, including glycerol (for cryopreservation of heterotrophs) and DMSO (Dimethyl sulfoxide, for cryopreservation of phototrophs).

Permitting and ethical compliance represent some of the most complex and time-intensive components of fieldwork planning and must be resolved before the collection of any material. Depending on the jurisdiction and sample type, obtaining the necessary permissions can require many months to over a year of advanced planning. Within the United States, land access and sampling regulations are generally defined by three major categories of land ownership: private, state or regional (e.g., municipal or county-managed), and federal (e.g., national parks or protected reserves). Each tier carries distinct permitting pathways, review timelines, and reporting requirements. Additionally, authorization to collect or transport biological material is further stratified by the type of sample. For example, CITES (Convention on International Trade in Endangered Species of Wild Fauna and Flora) permits are required for the collection or international transport of protected species (i.e., coral). This global treaty is designed to protect endangered plants and animals from international trade and promote legal and traceable trade of species (over 40,000 species are listed). While the USDA APHIS (Animal and Plant Health Inspection Service), under the United States government, regulates the importation of soils and other potential carriers of plant or animal pathogens to protect U.S. agriculture, it also requires permits for specific soil samples.

Samples associated with living hosts pose additional biosafety and pathogenicity risks and are therefore subject to heightened regulatory oversight, often requiring institutional biosafety review in addition to governmental approval. By contrast, environmental materials such as bulk water, eDNA, or marine sediment are typically regulated under broader, less restrictive import/export frameworks, provided they are not collected from designated protected or endangered habitats.

Private land is in many ways more straightforward as compared to other locations, as accessing it generally requires receiving permission from the relevant landowner and a mutually agreed-upon sample-usage scope signed and in writing. We provide an example of a very simple form of this agreement in the Handbook. Federal and State land in the U.S. typically have more formal permitting processes with longer lead times. However, for small amounts of material (e.g., a few milliliters), letters of exception can be issued to enable faster access. Outside of the U.S., there are generally analogs of these categories that each have their own usage designations.

Regardless of the permitting avenue and sample type, the downstream usage, relevant benefit sharing, and Nagoya protocol/High Seas Treaty compliance must be settled beforehand. The key point of consideration here is the degree to which a sample will be used for commercial or research purposes. Given the immense biotechnological value of environmental microbes, land managers are often within their rights to claim at least partial ownership of value derived from a downstream sample. The Nagoya Protocol attempts to govern this by mandating samples are available for research use and benefit sharing protocols are in-place. That said, benefit-sharing agreements with governments, while often well-intentioned, do not necessarily represent the interests of individual members of the society where samples are being collected. This is particularly the case for areas where Indigenous communities are the traditional custodians and long-stand stewards of the land and sea^24^.

Building on these considerations, and in addition to following local governmental policies and devising a clear Nagoya compliance strategy, the 2FP takes a sample-by-sample approach to permissions, ensuring that all outcomes will be open for research. Each sample we collect is annotated according to its potential downstream use. Commercial-use-available samples are annotated as such in case-by-case agreements set up with both relevant regulatory agencies as well as the landowner/land manager from where a sample is derived. We emphasize in the protocol the criticality of having local collaborators who live in and can speak directly to the interests of a community where sampling is ongoing and we provide recommendations for how to locate these individuals. We additionally provide a series of resources surrounding benefit sharing and ethical bioprospecting for use when designing a sampling protocol.

### Field sampling, processing, and transport

The middle sections of the handbook cover in-field activities, starting with field lab setup. In an ideal setting, a 2FP field lab is split into four different areas, staffed by dedicated team members. These include (1) the field sampling preparation area, where equipment is stored, cleaned, and restaged at the end of each day according to specific checklists, (2) a sample check-in area, where a sample is handed off from the field team to the lab team and receives a permanent, unique, pre-printed identifier generated with the CUAL-ID framework^25^, (3) a sample processing area, and (4) a sample preservation area, where processed (e.g., macerated) samples are inoculated into pre-filled and pre-labeled DNA storage buffer, glycerol, and DMSO tubes. These areas are separated in large part to maintain sterility – the preservation and processing areas, for example, are optimally in non-ventilated, low foot traffic regions where an open flame can be used for aseptic technique to minimize potential contamination. We additionally cover in this section waste disposal and guidelines for biosafety levels relevant to different sample types (e.g., requisite hood parameters for BSL-2 samples).

We delineate sample collection and downstream processing by both sample type and location. Generally speaking, sample types are classified as water, sediment, biomass (e.g., microbial mats), soils, animals (e.g., sponges, corals, insects), and plants, each of which requires slightly distinct protocols that are documented in both the handbook and the sampling slates. We collect samples in either conical tubes or Whirl-paks, though the latter are preferred to reduce the volume of material that must be carried or shipped (assuming water is filtered prior to shipment). All sample containers are prelabeled with waterproof pens (we reference the specific brand in the Resources table of this section) with a “Field ID;” a random, short identifier, usually encoding the sample type, day of sampling, and sample number (e.g., S1-1, S1-2, S1-3). Backpack coolers with ice packs are carried into the field, and the moment a sample is collected, it is cooled until it can be carried to the field lab for processing.

Sample check-in is the process of mapping a sample identifier to a hash that has been pre-generated by the CUAL-ID algorithm (referred to as the Two Frontiers ID, or TFID). Sample metadata (e.g., latitude and longitude, site and sample images, pH, geochemistry) is recorded, either being handwritten or recorded automatically with an custom application (e.g., qGIS), and a TFID is permanently assigned. A set number of TFIDs are generated and printed as part of the expedition planning process on cryolabels in rows of five. The first label is allocated to the sample collection container, the second to an intermediate processing vessel (e.g., a 50ml conical collecting filtered sample), and the remainder to DNA, glycerol, and DMSO tubes.

Processing protocols for each sample type are provided in the Handbook. For example, water samples are typically filtered via a vacuum pump to concentrate DNA, whereas corals are first homogenized with a sterile mortar and pestle before banking. These protocols are documented on both a slate as well as in the processing section of the handbook. Following the final stage of processing, samples are immediately aliquoted into a DNA preservative, glycerol, and DMSO and stored on dry ice. If dry ice is not available, we use pre-cooled phase change ice packs, ideally stored in a -40°C or -80°C freezer prior to departure for the field. If no other options are available, samples are kept on wet ice until they can be cryopreserved. At no point are samples allowed to increase in temperature following collection.

We additionally describe recommendations for the transport and shipping of samples. These include both safe packaging of biological material according to international shipping standards (e.g., double bagged with an absorbent material on the innermost bag) and strategies for maintaining sample viability during transit. Resources in this section include additional links out to permitting agencies and strategies for navigating customs effectively.

### Preservation strategy for sequencing and culturing compatibility

A primary goal of the 2FP workflow is to ensure that each collected sample remains viable for downstream applications, including high-throughput sequencing, cryopreservation, and culturing. To achieve this, 2FP employs a preservation system that utilizes aliquots stored in a DNA preservation buffer, glycerol, and DMSO. These methods help maintain sample integrity under varying field conditions, minimizes biomolecule and organism loss, and maximizes the recovery of large DNA and viable cells across diverse taxa for enrichment culture. Two cryoprotectants are used as their compound effectiveness varies between taxa. The preservation of environmental DNA is crucial for subsequent sequencing applications. However, conventional methods, such as ethanol fixation or refrigeration at 4 °C, often do not fully prevent enzymatic degradation, especially during transport or extended field deployment. DNA preservation buffers, such as DNA/RNA Shield, RNAlater, or equivalent formulations, chemically inactivate nucleases naturally present in the environment and can degrade a sample, preventing hydrolytic and oxidative damage even at ambient temperature. This eliminates the need for a continuous cold chain, which can be difficult to maintain in a field sampling environment, and yields higher-quality nucleic acids with fewer compositional biases than ethanol- or freezing-based storage.

In culture-based workflows, glycerol and DMSO serve as complementary cryoprotectants, preserving cellular viability during long-term storage and freeze–thaw cycles. Glycerol provides broad protection by reducing intracellular ice crystal formation, which can induce cell injury, as well as reducing osmotic shock across a wide range of bacteria and archaea. DMSO has been proven effective when preserving phototrophs and other microbial communities sensitive to cryopreservation, such as cyanobacteria and extremophiles, whose membrane structures render them more vulnerable to glycerol toxicity. By maintaining both glycerol- and DMSO-preserved aliquots, the 2FP workflow aims to maximize physiological and taxonomic diversity possible from a single environmental collection. Compared with freezing samples in bulk or storing them at 4 °C, this preservation system offers substantial logistical and scientific advantages. While freezing samples to −80 °C or below can be an effective way to preserve cells, it often compromises the integrity of the samples’ nucleic acid for long-term storage due to repeated freeze–thaw cycles, sublimation during long-term storage, or uneven temperature gradients within the sample matrix.

Additionally, prolonged freezing can reduce the likelihood of successful cell revival in microbial culturing, particularly for environmentally derived taxa. Although ethanol preservation and refrigeration are inexpensive and convenient, they provide minimal protection for live cells and can accelerate selective nucleic acid degradation in lysed microorganisms. This process disproportionately affects taxa with fragile cell envelopes or low GC-content genomes, thereby introducing compositional biases that can distort downstream molecular analyses, particularly in mixed or low-biomass environmental samples. In contrast, this schema stabilizes both genetic and physiological states at the point of collection, creating parallel archives that support metagenomic sequencing, culturing, and cryobanking without the need for redundant sampling or dedicated cold-chain infrastructure. This design ensures that each sample functions simultaneously as a genomic record and a living resource—enabling direct linkage between community composition, functional potential, and experimentally validated phenotypes.

### Post-sampling debrief and process development

The final Handbook section covers the sampling debrief. This is an all hands, retrospective meeting where challenges are described and discussed in a structured format. The aim in this meeting, which ideally would happen within two weeks of sampling, is to not only identify problems, but also construct solutions and their associated, specific implementation strategies. This plan-execute-debrief cycle is how we built this resource, which is at present on version 11.0.

### Supporting and streamlining sampling with 2FP OpenTools

We also provide a suite of open hardware and software that is either referenced in or generally relevant to the operations described in the Handbook. This toolkit is referred to as the 2FP OpenTools, and it is currently reflected in the contents of numerous GitHub repositories, all summarized on the Handbook website (Table 2). They include both one-off kits and protocols we have constructed for different community science sampling projects, software packages developed for analysis of metagenomic and bacterial isolate data, and protocol slates. The toolkit also contains an expedition template repository, which includes our standard, version-controlled, Excel-compatible template forms to be used before, during, and after sampling for tasks ranging from budgeting to metadata entry and team debriefs. The hardware included in the 2FP OpenTools ranges from simple 3D printed racks and mounts for a field lab to a low-cost spectrophotometer. We additionally detail a low-cost 3D printed microscope that we have used in the field numerous times for evaluating organismal morphology and diversity on-site. All equipment not originally designed by the 2FP team are appropriately referenced in its repository, giving credit to the original creator.

**Table 2.** Hardware in the 2FP OpenTools collection.

| **Tool** | **Summary** |
| --- | --- |
| **Dremelfuge** | A portable centrifuge adapter compatible with Dremel rotary tools for rapid field centrifugation. |
| **3D-printed Tools** | Includes eyeglasses mount for divemasks, laser cut racks for efficient organization of cryovials, various bottle holders and adapters for field tools (e.g., a cuvette holder). |
| **PUMA Scope** | Compact, field-deployable microscope with FreeCAD design files for microscopy under rugged conditions. |
| **UVspec** | Open ultraviolet spectrometry hardware for quantifying absorbance or fluorescence in the field. |
| **OD600** | DIY optical density meter for microbial growth or turbidity measurements at 600 nm. |
| **Open Colorimeter** | Portable, open-source colorimeter for measuring concentration and absorbance in field samples. |

**Table 3.** Software in the 2FP OpenTools collection.

| **Tool** | **Summary** |
| --- | --- |
| **XTree** | Ultra-fast, pan-domain alignment and taxonomy software for metagenomic and genomic datasets. |
| **MAGUS** | Comprehensive assembly, binning, and annotation pipeline for multi-domain environmental genomics. |

## Discussion

The 2FP Handbook and OpenTools Resource is designed for use by the scientific community to assay the extraordinary diversity of microbial life on the planet while also considering the strategic, personnel, and ethical considerations of large complex sampling operations. The resource comprises the 2FP Handbook itself, its accompanying in-field sampling slates, and the complete set of public repositories and toolkits used and maintained by The Two Frontiers Project (https://two-frontiers-project.github.io/#/).

We anticipate that we will continue to improve and integrate numerous additions to the Handbook and Toolkit as we continue to use this workflow. These include our in-field extraction, library preparation, and sequencing methods as well as additional protocols that we use after returning from the field. We also will provide tools relevant for on-site culturing and media design (e.g., low cost field incubator setups). Finally, we aim to develop a separate set of web portal-based workflows regarding permitting, sampling, and open science. This will include web portal-based tools for assisting teams in navigating the permitting and experimental design process, providing an interface to plan sequencing approaches, determine requisite sample sizes, and plan logistics.

Overall, we hope that the tools here provide a centralized basis for conducting and executing sampling campaigns for environmental microbiology. We are confident that unlocking this diversity in a sustainable and ethical manner will have dramatic implications for human and planetary health. We welcome feedback from the scientific community and look forward to further developing this resource.

## Methods

### Design of the 2FP Handbook and OpenTools Resource

The Two Frontiers Project was founded in the fall of 2022 and became an independent, 501(c)(3) non-profit in mid-2024. Since our first expedition, in September 2022, we have been developing and optimizing protocols for field microbiology that met the requisite standards for downstream cryopreservation, sequencing, and culturing. We have developed these tools over 12+ large-scale (multi-day or in remote locales) expeditions in the intervening time, as well as numerous smaller sampling campaigns and two community science initiatives that spanned the United States. The Resource described in this manuscript represents the aggregation of that effort. The Handbook itself is currently on Version 11.0

## Supporting information

Supplementary Text S1. & Table S1 Protocol Iteration and Stabilization

Supplementary Table S2. Field Expeditions Contributing to Protocol Development and Validation

Supplementary File S3 Example expedition planning and execution package (completed template) V11

## Data Availability

All resources described in this study are available at https://two-frontiers-project.github.io/.

---

Supplementary Text S1. Protocol Iteration and Stabilization

Supplementary Table S1. Early protocol versions

Supplementary Table S2. Field Expeditions Contributing to Protocol Development and Validation

Supplementary File S3: Example expedition planning and execution package (completed template)

## References

1. Whitman, W.B., Coleman, D.C., and Wiebe, W.J. (1998). Prokaryotes: the unseen majority. Proc. Natl. Acad. Sci. U. S. A. 95, 6578–6583.

2. Zimmerman, S., Tierney, B.T., Patel, C.J., and Kostic, A.D. (2023). Quantifying shared and unique gene content across 17 microbial ecosystems. mSystems 8, e0011823.

3. Coelho, L.P., Alves, R., Del Río, Á.R., Myers, P.N., Cantalapiedra, C.P., Giner-Lamia, J., Schmidt, T.S., Mende, D.R., Orakov, A., Letunic, I., et al. (2022). Towards the biogeography of prokaryotic genes. Nature 601, 252–256.

4. Schultz, J., Modolon, F., Peixoto, R.S., and Rosado, A.S. (2023). Shedding light on the composition of extreme microbial dark matter: alternative approaches for culturing extremophiles. Front. Microbiol. 14, 1167718.

5. Rinke, C., Schwientek, P., Sczyrba, A., Ivanova, N.N., Anderson, I.J., Cheng, J.-F., Darling, A., Malfatti, S., Swan, B.K., Gies, E.A., et al. (2013). Insights into the phylogeny and coding potential of microbial dark matter. Nature 499, 431–437.

6. Mayorga-Ramos, A., Zúñiga-Miranda, J., Carrera-Pacheco, S.E., Barba-Ostria, C., and Guamán, L.P. (2023). CRISPR-Cas-based antimicrobials: Design, challenges, and bacterial mechanisms of resistance. ACS Infect. Dis. 9, 1283–1302.

7. Rugarabamu, S., and Mwanyika, G. (2025). Biotechnological innovations to combat antimicrobial resistance and advance global health equity. Bacteria 4, 46.

8. Santos-Beneit, F. (2024). What is the role of microbial biotechnology and genetic engineering in medicine? Microbiologyopen 13, e1406.

9. Garcias-Bonet, N., Roik, A., Tierney, B., García, F.C., Villela, H.D.M., Dungan, A.M., Quigley, K.M., Sweet, M., Berg, G., Gram, L., et al. (2024). Horizon scanning the application of probiotics for wildlife. Trends Microbiol. 32, 252–269.

10. Peixoto, R., Voolstra, C.R., Stein, L.Y., Hugenholtz, P., Salles, J.F., Amin, S.A., Häggblom, M., Gregory, A., Makhalanyane, T.P., Wang, F., et al. (2024). Microbial solutions must be deployed against climate catastrophe. Nat. Commun. 15, 9637.

11. Gilbert, J.A., Peixoto, R.S., Scholz, A.H., Dominguez Bello, M.G., Korsten, L., Berg, G., Singh, B., Boetius, A., Wang, F., Greening, C., et al. (2025). Launching the IUCN Microbial Conservation Specialist Group as a global safeguard for microbial biodiversity. Nat. Microbiol. 10, 2359–2360.

12. Modolon, F., Schultz, J., Duarte, G., Vilela, C.L.S., Thomas, T., and Peixoto, R.S. (2023). In situ devices can culture the microbial dark matter of corals. iScience 26, 108374.

13. Matar, G., and Bilen, M. (2022). Culturomics, a potential approach paving the way toward bacteriotherapy. Curr. Opin. Microbiol. 69, 102194.

14. Wynberg, R. (2023). Biopiracy: Crying wolf or a lever for equity and conservation? Res. Policy 52, 104674.

15. Heinz, E., Holt, K.E., Meehan, C.J., and Sheppard, S.K. (2021). Addressing parachute research and removing barriers for LMIC researchers in Microbial Genomics. Microb. Genom. 7. 10.1099/mgen.0.000722.

16. Macilwain, C. (1998). When rhetoric hits reality in debate on bioprospecting. Nature 392, 535–537, 539–540.

17. Beato, M.S., and Veneroso, V. (2023). The Nagoya Protocol on access and benefit sharing: The neglected issue of animal health. Front. Microbiol. 14. 10.3389/fmicb.2023.1124120.

18. The Nagoya Protocol on Access and Benefit-sharing (2025). Secretariat of the Convention on Biological Diversity. https://www.google.com/url?q= https://www.cbd.int/abs/default.shtml&sa=D&source=docs&ust=1761330062440792&usg=AOvVaw0Gh-befOxNeGeDthzKEF3_.

19. https://www.un.org/bbnjagreement/sites/default/files/2024-08/Text.

20. Buck, M., and Hamilton, C. (2011). The Nagoya Protocol on access to Genetic Resources and the Fair and Equitable Sharing of Benefits Arising from their Utilization to the Convention on Biological Diversity: The Nagoya protocol. Rev. Eur. Community Int. Environ. Law 20, 47–61.

21. Rourke, M.F. Access and benefit-sharing in practice: non-commercial research scientists face legal obstacles to accessing genetic resources. https://www.google.com/url?q=https://www.sciencepolicyjournal.org/uploads/5/4/3/4/5434385/rourke.pdf&sa=D&source=docs&ust=1761330062438892&usg=AOvVaw3K4tDvyVeknVwPHep6AJ3P.

22. Wood-Charlson, E.M., Anubhav Auberry, D., Blanco, H., Borkum, M.I., Corilo, Y.E., Davenport, K.W., Deshpande, S., Devarakonda, R., Drake, M., et al. (2020). The National Microbiome Data Collaborative: enabling microbiome science. Nat. Rev. Microbiol. 18, 313–314.

23. Kelliher, J.M., Mirzayi, C., Bordenstein, S.R., Oliver, A., Kellogg, C.A., Hatcher, E.L., Berg, M., Baldrian, P., Aljumaah, M., Miller, C.M.L., et al. (2025). STREAMS guidelines: standards for technical reporting in environmental and host-associated microbiome studies. Nat. Microbiol. 10, 3059–3068.

24. Liggins, L., Hudson, M., and Anderson, J. (2021). Creating space for Indigenous perspectives on access and benefit-sharing: Encouraging researcher use of the Local Contexts Notices. Mol. Ecol. 30, 2477–2482.

25. Chase, J.H., Bolyen, E., Rideout, J.R., and Caporaso, J.G. (2016). Cual-id: Globally unique, correctable, and human-friendly sample identifiers for comparative omics studies. mSystems 1. 10.1128/mSystems.00010-15.

