## Supplementary Text S1. & Table S1 Protocol Iteration and Stabilization for "The Two Frontiers Project Field Handbook and OpenTools: Standardizing microbial fieldwork for biobank-scale sequencing and culturomics"

*Supplementary Text S1. Protocol Iteration and Stabilization*

Protocols used in this study were developed and refined iteratively over a three-year period of field sampling, laboratory processing, and sample transportation across diverse terrestrial and aquatic environments (Supplementary Table S2). Revisions were implemented in response to operational constraints encountered during expeditions, with the goal of improving sampling consistency, contamination control, preservation outcomes, metadata completeness, and chain-of-custody traceability. Early protocol versions focused on establishing minimum viable workflows for metadata capture and sample preservation that could be executed reliably in field settings. Subsequent revisions standardized sampling order across environments to reduce cross-contamination, introduced SCUBA-specific guidance for submerged sampling, and unified processing steps prior to preservation to ensure consistency across sample types. Later versions formalized temperature targets and monitoring during transport, clarified the handling of negative controls and blanks, and standardized identifier linkage across physical samples and digital records.
The final protocol release (v11.0) consolidated these components into workflow-aligned field slates to improve usability and protocol adherence during multi-site campaigns. Only revisions affecting core operational outcomes were considered substantive and are summarized below. Minor editorial updates, formatting changes, and non-operational clarifications were excluded. All protocol versions are archived in a public, version-controlled repository (https://two-frontiers-project.github.io/).

Supplementary Table S1.

| Protocol Domain | Version | Year Implemented | Substantive Change Implemented | Operational Rationale | Outcome / Impact |
|---|---|---|---|---|---|
| Field collection | v1 | 09/2022 | Established baseline field collection procedures for water, sediment, and biomass sampling. | Enable initial multi-site sampling across environments. | Enabled early comparative sampling campaigns. |
| Field preservation | v2 | 02/2023 | Introduced parallel preservation using DNA preservation buffer and cryoprotectants. | Maintain DNA integrity and cell viability under field conditions. | Improved downstream sequencing and culturing success. |
| Standardized metadata capture | v3 | 03/2023 | Standardized minimum required metadata fields across all sample types | Reduce missing or inconsistent metadata across campaigns. | Increased metadata completeness and cross-site comparability. |
| Field sampling order | v4 | 09/2023 | Defined fixed sampling order | Minimize cross-contamination between sample types. | Reduced contamination risk in low-biomass samples. |
| Underwater sampling | v5 | 03/2024 | Introduced SCUBA-specific contamination controls and buoyancy guidance. Handling issues under deep dives (>60m in depth). | Address constraints unique to submerged sampling. | Improved sample integrity during underwater collection. |
| Identifier linkage across preservation types | v6 | 04/2024 | Standardized linkage between Field IDs, CIDs, and TFIDs across all aliquots. | Ensure traceability across physical samples and digital records. | Reduced accessioning and labeling errors. Internal lab processing and organization simplified. |

Supplementary Table S1. continued

| Protocol Domain | Version | Year Implemented | Substantive Change Implemented | Operational Rationale | Outcome / Impact |
|---|---|---|---|---|---|
| Sample processing | v7 | 08/2025 | Unified pre-preservation processing workflow across sample types. Removed steps to subset samples. | Ensure consistency prior to aliquoting and preservation. | Improved reproducibility of downstream analyses. |
| Temperature control and transport | v8 | 10/2025 | Defined temperature targets and monitoring requirements for transport, especially with cryopreserved samples. | Reduce degradation during domestic and international shipment. | Improved sample stability during transit. |
| Bulk water collection | v9 | 03/2025 | Formalized bulk water collection volumes, containers, and filtration strategy. | Enables scalable water sampling without compromising transport; increases recovery of microbial biomass from low-biomass environments. | Improves consistency across high-volume water samples; increases DNA yield and cultured biomass. |
| Kit-based sampling strategy | v10 | 07/2025 | Standardized modular field sampling kits tailored by environment and sample type. | Reduce field variability and preparation errors | Increased operational efficiency and protocol adherence. |
| Workflow integration | v11 | 11/2025 | Consolidated protocols into workflow-aligned field slates. | Improve usability and adherence in field and laboratory settings. | Increased consistency across teams and campaigns. |
