## Supplementary Table S2. Field Expeditions Contributing to Protocol Development and Validation for "The Two Frontiers Project Field Handbook and OpenTools: Standardizing microbial fieldwork for biobank-scale sequencing and culturomics"

<u>Description:</u> Summary of field expeditions conducted during the study period in which the 2FP Field Handbook protocols were applied, evaluated, and refined. Expeditions are listed chronologically and include both terrestrial and marine sampling campaigns.

| Expedition | Location | Country | Environment Type | Timeline |
|---|---|---|---|---|
| Carbon1 | Vulcano | Italy | Terrestrial geothermal | September 2022 |
| Coral1 | Red Sea | Saudi Arabia | Marine / coral-associated | December 2022 |
| Carbon2 | Colorado (San Juan National Forest) | USA | Terrestrial geothermal | February 2023 |
| Carbon3 | La Terra Fumante (“The Smoking Lands”) | Italy | Terrestrial geothermal | May 2024 |
| Carbon4 | Shikinejima | Japan | Terrestrial geothermal / coastal | August 2024 |
| Carbon5 | Ambitle Island | Papua New Guinea | Volcanic / coastal geothermal | October 2024 |
| Crops1 | Mojave Preserve | USA | Terrestrial | December 2024 |
| Carbon6 | Colorado | USA | Terrestrial geothermal | February 2025 |
| Carbon7 | Vulcano | Italy | Terrestrial geothermal | July 2025 |
| Methane1 | Scoglio d’Africa | Italy | Marine methane seep | July 2025 |
| Carbon8 | Colorado | USA | Terrestrial geothermal | September 2025 |
| Methane2 | Buzău Land | Romania | Terrestrial methane seep / mud volcanoes | December 2025 |
