## Supplementary File S3 Example expedition planning and execution package (completed template) V11 for "The Two Frontiers Project Field Handbook and OpenTools: Standardizing microbial fieldwork for biobank-scale sequencing and culturomics": 2fp_research_proposal_carbon9.docx

Expedition Name: Spence Hot Springs

Project: The Two Frontiers Project (2FP)
 Program Area: Carbon / Extreme Terrestrial Environments
 Location: Spence Hot Springs, Santa Fe National Forest / Valles Caldera region, New Mexico, USA

Purpose of Expedition:
This expedition aims to characterize microbial communities associated with geothermal hot spring environments in a temperate continental setting. Spence Hot Springs was selected as an accessible, low-temperature geothermal site to support pilot sampling of microbial mats, sediments, and spring water.

Primary Objectives:

- Collect microbial mat, sediment, and water samples from a geothermal spring system
- Record standardized environmental metadata for integration into the 2FP Living Database
- Generate baseline data to inform future sampling within the Valles Caldera region

Dates:

Field Sampling: May 18–19, 2025

Study Site Description:

Spence Hot Springs is a series of geothermal pools located along the Jemez River within the Santa Fe National Forest, adjacent to the Valles Caldera region. The site is characterized by warm spring outflows, mineral-rich sediments, and visible microbial mats. Water temperatures are moderate relative to other geothermal systems, making the site suitable for pilot sampling and method validation.

Sampling Overview:
 Sampling focused on three primary substrates:

- Surface microbial mats associated with spring outflow areas
- Sediment collected from shallow pool margins
- Spring water collected upstream and downstream of visible geothermal influence

Environmental metadata collected included GPS coordinates, water temperature, pH, visual site observations, and weather conditions at the time of sampling.

Permitting and Access:
Sampling will be conducted under permissions consistent with Santa Fe National Forest regulations for non-invasive scientific sampling. No mechanized equipment or permanent site alteration was performed. All sampling followed Leave No Trace principles.

Logistics Summary:
The site will be accessed via established hiking trails. All equipment to be transported by hand. Samples to be temporarily stored in insulated containers and transported to the laboratory under controlled temperature conditions on the same day.

Projected Outcomes:

- Successful collection of paired biological samples and metadata
- Identification of site-specific considerations for future geothermal sampling campaigns
