## Supplementary File S3 Example expedition planning and execution package (completed template) V11 for "The Two Frontiers Project Field Handbook and OpenTools: Standardizing microbial fieldwork for biobank-scale sequencing and culturomics": Carbon9_ Spence.pptx

### Slide 1
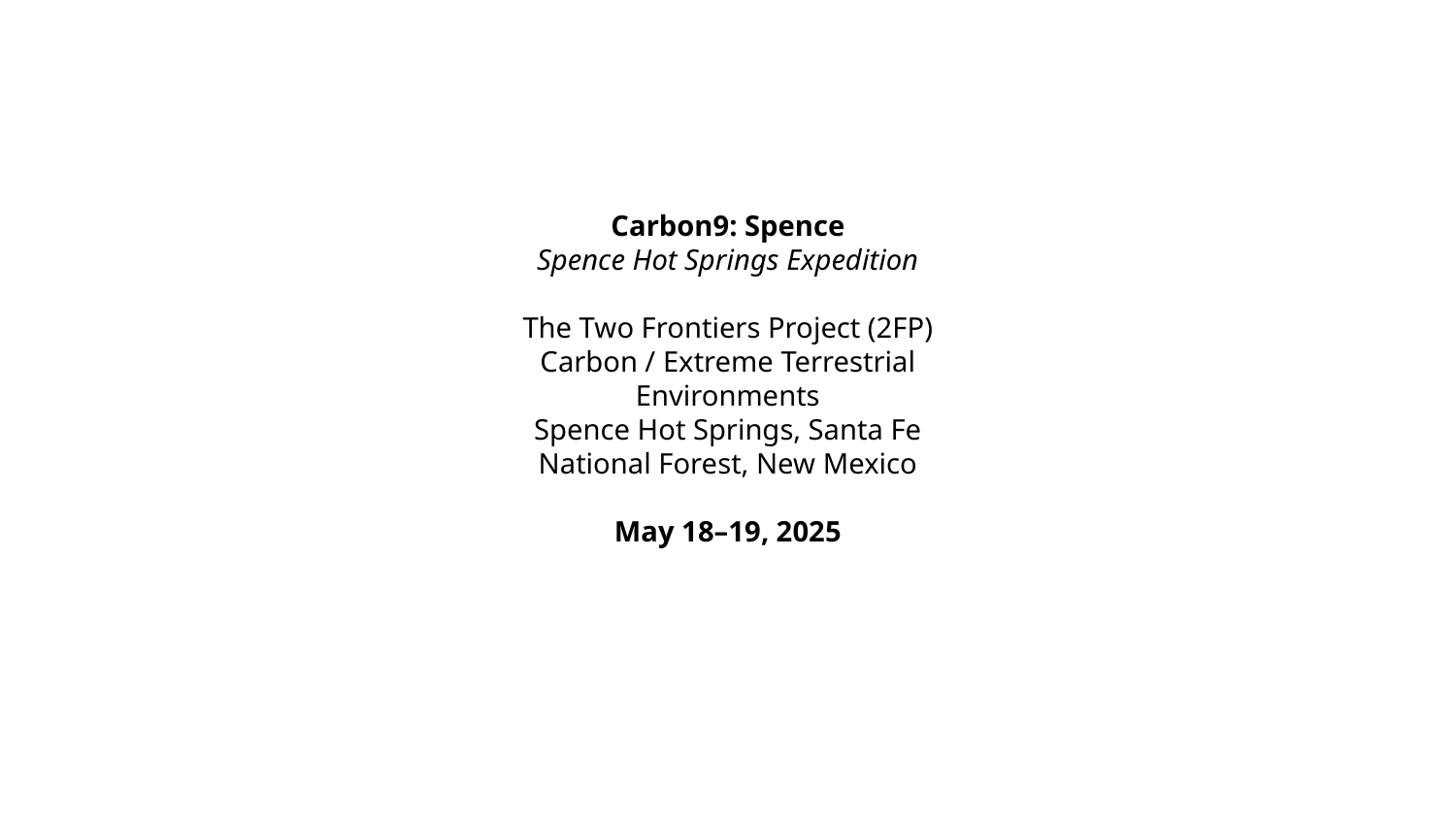

Carbon9: Spence
Spence Hot Springs Expedition
The Two Frontiers Project (2FP)
Carbon / Extreme Terrestrial Environments
Spence Hot Springs, Santa Fe National Forest, New Mexico
May 18–19, 2025

### Slide 2
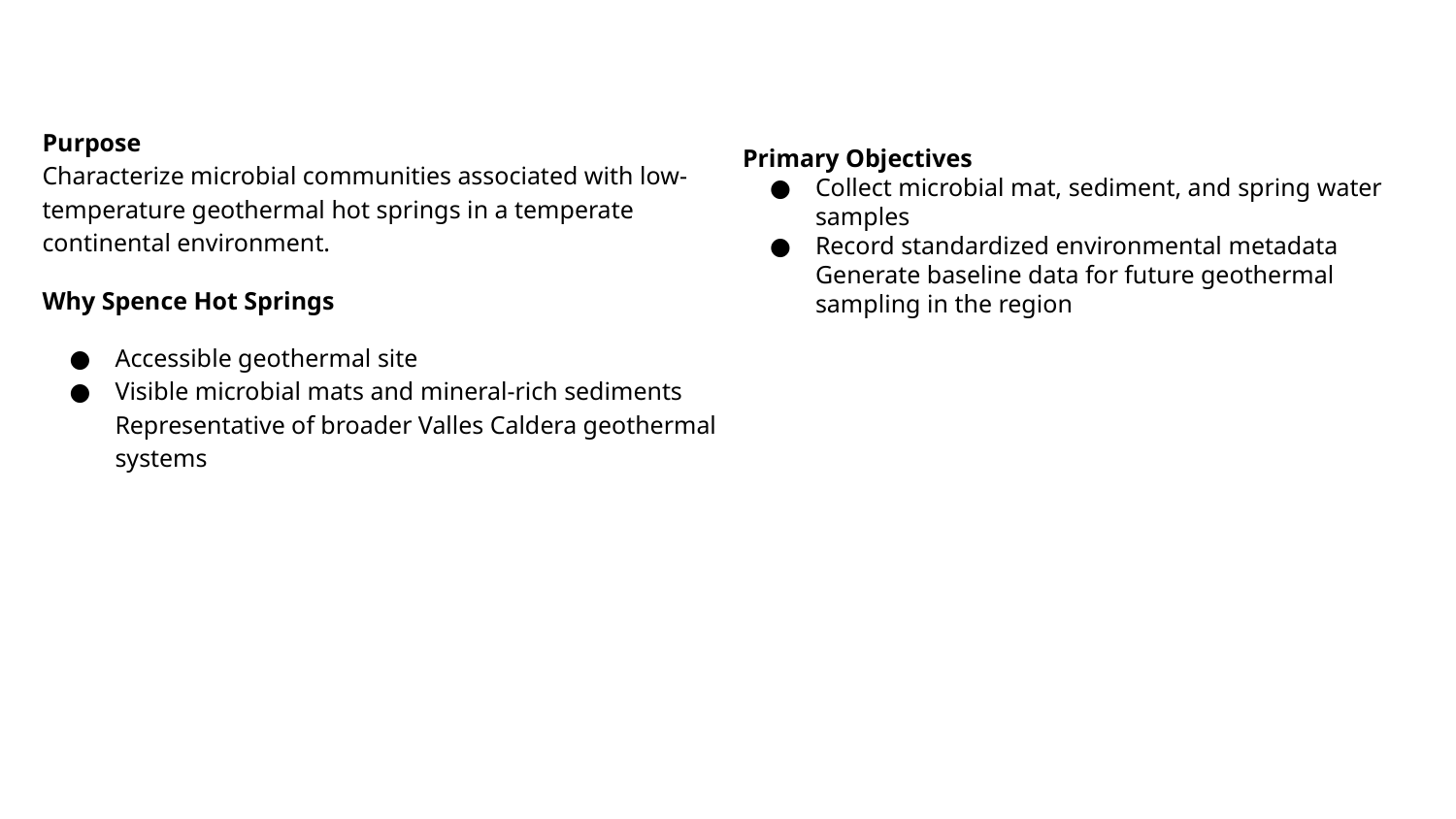

Purpose
Characterize microbial communities associated with low-temperature geothermal hot springs in a temperate continental environment.
Why Spence Hot Springs
Accessible geothermal site
Visible microbial mats and mineral-rich sediments
Representative of broader Valles Caldera geothermal systems
Primary Objectives
Collect microbial mat, sediment, and spring water samples
Record standardized environmental metadata
Generate baseline data for future geothermal sampling in the region

### Slide 3
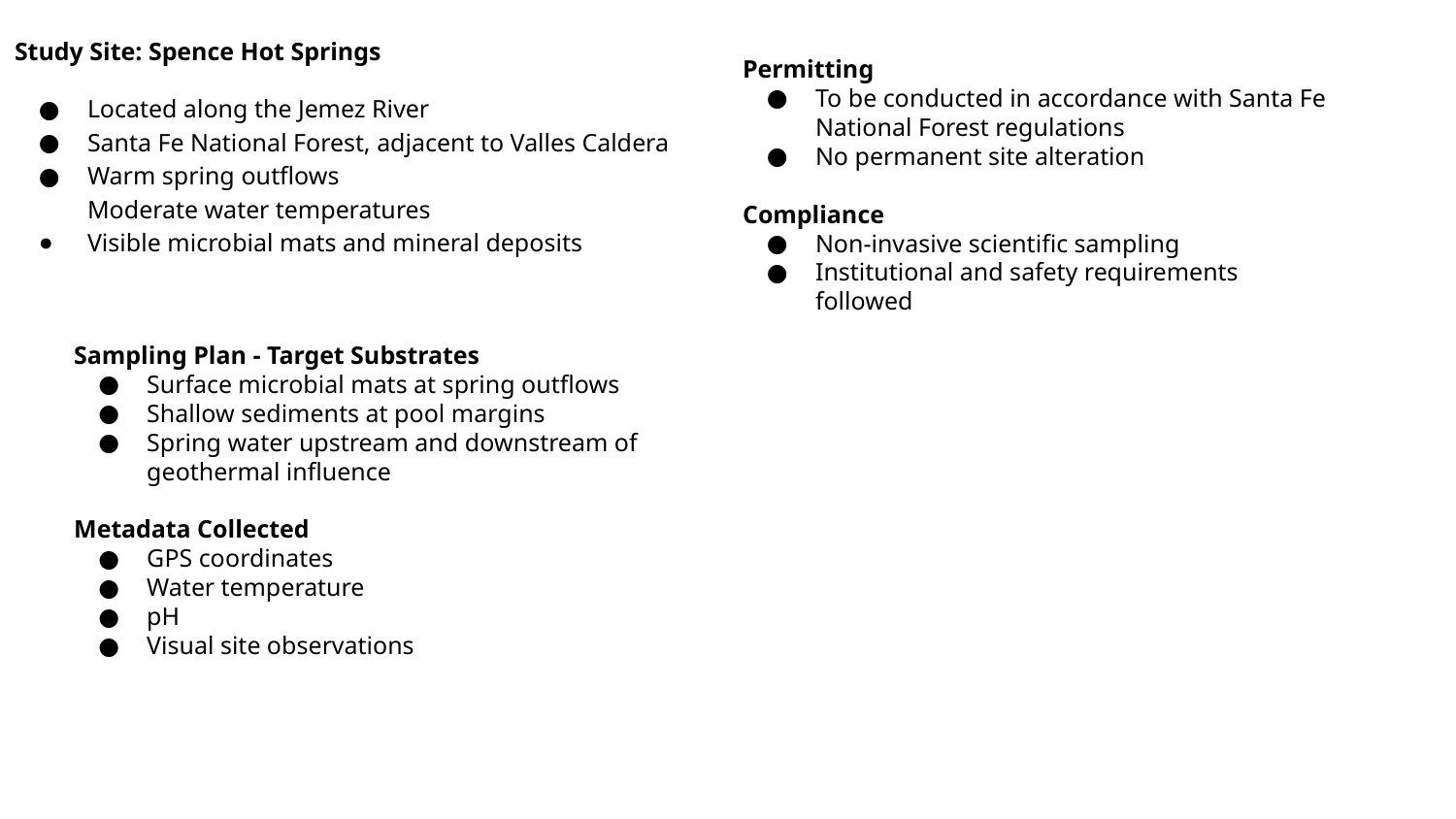

Study Site: Spence Hot Springs
Located along the Jemez River
Santa Fe National Forest, adjacent to Valles Caldera
Warm spring outflows
Moderate water temperatures
Visible microbial mats and mineral deposits
Permitting
To be conducted in accordance with Santa Fe National Forest regulations
No permanent site alteration
Compliance
Non-invasive scientific sampling
Institutional and safety requirements followed
Sampling Plan - Target Substrates
Surface microbial mats at spring outflows
Shallow sediments at pool margins
Spring water upstream and downstream of geothermal influence
Metadata Collected
GPS coordinates
Water temperature
pH
Visual site observations
