## Supplementary File S3 Example expedition planning and execution package (completed template) V11 for "The Two Frontiers Project Field Handbook and OpenTools: Standardizing microbial fieldwork for biobank-scale sequencing and culturomics": README_01.docx

Folder: 01_BACKGROUND_RESEARCH

Expedition: [Expedition Name]

This folder contains optional background and preparatory materials assembled during the early planning stages of a 2FP expedition. These materials provide scientific, logistical, and contextual support for expedition design but are not required for field execution.

Files in this folder may include, but are not limited to:

- Literature and reference materials relevant to target environments, organisms, or research questions
- Early planning documents, including candidate dates, locations, and site assessments
- Research proposals, concept notes, or funder briefing materials
- Contact lists and correspondence used during preliminary coordination with collaborators, partners, or site managers
- Presentation materials (e.g., slides) developed for internal planning or external discussions

To preserve consistency and reproducibility across expeditions, the following should not be stored in this folder:

- Final sampling protocols or SOPs (see 04_PROTOCOLS_EXPERIMENTS)
- Field maps, GIS layers, or site coordinates (see 02_FIELD)
- Permits or regulatory documents (see 03_PERMITTING)
- Operational logistics, inventories, or travel plans (see expedition planning files)
