## Supplementary File S3 Example expedition planning and execution package (completed template) V11 for "The Two Frontiers Project Field Handbook and OpenTools: Standardizing microbial fieldwork for biobank-scale sequencing and culturomics": README_02.docx

```
Folder: 02_FIELD (MAPS, GIS)
 Project: The Two Frontiers Project (2FP)
 Expedition: [Expedition Name]
This folder contains field-facing spatial materials used during site selection, navigation, and sampling execution.
Typical contents include:
```

- Regional and site-level maps (PDF or image format)
- GIS layers and coordinate files
  Route planning maps and access paths
- Screenshots or exports shared by local collaborators
  Field map annotations or quick-reference visuals

This folder should contain finalized field maps only. Background literature maps should be stored in 01_BACKGROUND_RESEARCH.
