## Supplementary File S3 Example expedition planning and execution package (completed template) V11 for "The Two Frontiers Project Field Handbook and OpenTools: Standardizing microbial fieldwork for biobank-scale sequencing and culturomics": Permitting Example ΓÇö Santa Fe National Forest.docx

Expedition: Spence Hot Springs

The Two Frontiers Project (2FP)

Sampling for this expedition will be conducted within lands managed by the Santa Fe National Forest. Activities are designed to fall within the scope of non-invasive, low-impact scientific observation and sample collection, consistent with U.S. Forest Service regulations.

Planned field activities include manual collection of small-volume water samples, surface microbial mats, and shallow sediments using hand tools only. No mechanized equipment, excavation, drilling, or permanent site modification will be performed.

Permitting and access requirements were reviewed in advance with Santa Fe National Forest guidance for research activities. Sampling is conducted in accordance with applicable Forest Service policies governing special uses and research on public lands. Where required, notification or authorization is obtained through the appropriate Forest Service office prior to field activities.

All sampling locations are accessed via established trails or previously disturbed areas. Field teams follow Leave No Trace principles, including minimizing disturbance, restoring sampling sites to their original condition where applicable, and packing out all materials.

No collection of protected species is planned, and no samples are taken from culturally or environmentally sensitive areas. Sampling volumes are limited to the minimum necessary to meet scientific objectives.

This permitting framework serves as an example of how low-risk pilot sampling campaigns may be conducted within U.S. National Forest lands, and does not replace site-specific permitting requirements for future expeditions.
