## Supplementary File S3 Example expedition planning and execution package (completed template) V11 for "The Two Frontiers Project Field Handbook and OpenTools: Standardizing microbial fieldwork for biobank-scale sequencing and culturomics": README_03.docx

```
Folder: 03_PERMITTING
 Project: The Two Frontiers Project (2FP)
 Expedition: [Expedition Name]
This folder contains all regulatory, legal, and access documentation required to conduct the expedition.
Typical contents include:
```

- Permits and access agreements
- Memoranda of understanding or cooperation agreements
- Letters of authorization from site managers or authorities
- Institutional forms (e.g., equipment declarations)
- Email correspondence related to permitting
- Drafts and finalized versions of submitted documents

Only finalized or actively relevant permitting materials should be stored here.
