## Supplementary File S3 Example expedition planning and execution package (completed template) V11 for "The Two Frontiers Project Field Handbook and OpenTools: Standardizing microbial fieldwork for biobank-scale sequencing and culturomics": README_04.docx

```
Folder: 04_PROTOCOLS_EXPERIMENTS
 Project: The Two Frontiers Project (2FP)
 Expedition: [Expedition Name]
This folder contains scientific protocols and reference materials used during field sampling and experiments.
Typical contents include:
```

- Sampling and processing protocols
- Instrument or sensor operating procedures
- Relevant sections of the 2FP Field Handbook
- Reference diagrams or figures used in the field

Draft planning documents should be stored in 01_BACKGROUND_RESEARCH.

See full handbook in GitHub.
