## Supplementary File S3 Example expedition planning and execution package (completed template) V11 for "The Two Frontiers Project Field Handbook and OpenTools: Standardizing microbial fieldwork for biobank-scale sequencing and culturomics": README_05.docx

```
Folder: 05_PHOTOS
 Project: The Two Frontiers Project (2FP)
 Expedition: [Expedition Name]
This folder contains photographic documentation collected during the expedition.
Typical contents include:
```

- Site photos
- Sample photos
- Lab notebook or workflow photos
- Drone imagery
- Press or outreach photos
- Other contextual images relevant to the expedition

Subfolders are used to separate photo types. Image metadata and naming conventions should follow 2FP photo guidelines where applicable.
