## Supplementary File S3 Example expedition planning and execution package (completed template) V11 for "The Two Frontiers Project Field Handbook and OpenTools: Standardizing microbial fieldwork for biobank-scale sequencing and culturomics": README_06.docx

```
Folder: 06_BILLING_RECEIPTS
 Project: The Two Frontiers Project (2FP)
 Expedition: [Expedition Name]
This folder contains financial documentation associated with expedition expenses.
Typical contents include:
```

- Travel receipts (flights, trains, buses)
- Lodging invoices
- Equipment or supply receipts
- Other reimbursable expense documentation

Receipts should be organized into subfolders where possible. No analysis or summaries should be stored here.
