## Supplementary File S3 Example expedition planning and execution package (completed template) V11 for "The Two Frontiers Project Field Handbook and OpenTools: Standardizing microbial fieldwork for biobank-scale sequencing and culturomics": Round trip to Albuquerque _ Google Flights.pdf

Share

### Boston ⇄ Albuquerque

**$354**

Round trip · Economy (include Basic) · 1 passenger

Lowest total price

#### Selected flights

##### Departing flight · Thu, May 7

jetBlue

**2:55 PM**
Boston Logan International Airport (BOS)

Travel time: 1 hr 18 min

**4:13 PM**
John F. Kennedy International Airport (JFK)

JetBlue B6 517 · Economy
Often delayed by 30+ min

- Above average legroom (32 in)
- Free Wi-Fi
- In-seat power & USB outlets
- Live TV

3 hr 16 min layover · New York (JFK)

jetBlue

**7:29 PM**
John F. Kennedy International Airport (JFK)

Travel time: 4 hr 50 min

**10:19 PM**
Albuquerque International Sunport (ABQ)

JetBlue B6 65 · Economy · Airbus A320

- Above average legroom (32 in)
- Free Wi-Fi
- In-seat power & USB outlets
- Live TV
- Emissions estimate: 260 kg CO2e
- Contrail warming potential: High

##### Returning flight · Thu, May 14

Travel time: 4 hr 11 min · Overnight ⚠

5:25 AM+1
John F. Kennedy International Airport (JFK)

JetBlue B6 66 · Economy · Airbus A320

- Above average legroom (32 in)
- Free Wi-Fi
- In-seat power & USB outlets
- Live TV
- Emissions estimate: 265 kg CO2e
- Contrail warming potential: Medium ⓘ

1 hr 5 min layover · New York (JFK)

6:30 AM+1
John F. Kennedy International Airport (JFK)

Travel time: 1 hr 11 min

7:41 AM+1
Boston Logan International Airport (BOS)

JetBlue B6 118 · Economy

- Above average legroom (32 in)
- Free Wi-Fi
- In-seat power & USB outlets
- Live TV

1 free carry-on    1st checked bag available for a fee

Baggage conditions apply to your entire trip. Bag fees may be higher at the airport. JetBlue bag policy

#### Booking options

How options are ranked ⓘ

| | | |
|---|---|---|
| Book with Orbitz | $354 | Continue |
| Book with Travelocity | $354 | Continue |

Prices include required taxes + fees for 1 adult. Optional charges and bag fees may apply.

Displayed currencies may differ from the currencies used to purchase flights.
Learn more
