## Supplementary File S3 Example expedition planning and execution package (completed template) V11 for "The Two Frontiers Project Field Handbook and OpenTools: Standardizing microbial fieldwork for biobank-scale sequencing and culturomics": README_07.docx

```
Folder: 07_FORMS
 Project: The Two Frontiers Project (2FP)
 Expedition: [Expedition Name]
This folder contains completed or reference forms required for expedition participation and compliance.
Typical contents include:
```

- Field waivers
- Safety or liability forms
- Diving or activity-specific forms
  Institutional or partner-required documentation

Blank templates should be stored elsewhere; this folder is intended for expedition-specific forms.
