## Supplementary File S3 Example expedition planning and execution package (completed template) V11 for "The Two Frontiers Project Field Handbook and OpenTools: Standardizing microbial fieldwork for biobank-scale sequencing and culturomics": The Two Frontiers Project ΓÇô Expedition Field Waiver - Google Forms.pdf

### The Two Frontiers Project – Expedition Field Waiver

Please complete this form before participating in any 2FP fieldwork. It outlines important safety, liability, and consent information for your protection and ours. All responses are confidential and securely stored.

1. Full Name

2. Date

   *Example: January 7, 2019*

3. Address

4. Phone Number (with country code)

5. Email

6. Date of Birth

_______________________________

*Example: January 7, 2019*

7. **Emergency Contact:** Full Name

_______________________________

8. **Emergency Contact:** What is your relationship to your emergency contact?

   *Mark only one oval.*

   - Parent/Guardian
   - Sibling
   - Spouse/Partner
   - Family Member
   - Friend
   - Colleague
   - Other: _______________________________

9. **Emergency Contact:** Phone Number

_______________________________

#### ASSUMPTION OF RISK

Participation in field research and expeditionary work may involve a variety of risks, including environmental, physical, medical, and travel-related hazards. This section outlines the types of risks you may encounter while involved in 2FP activities. By initialing below, you acknowledge that you understand these risks and voluntarily accept them as a condition of participation If you have any questions, please email **

We ask the participant to acknowledge that involvement with 2FP, including but not limited to participation in research activities, fieldwork, workshops, conferences, or other engagements, may involve risks. By participating in 2FP fieldwork, expeditions, or research activities, you may be exposed to a range of risks, including but not limited to:

10. **Environmental & Physical Hazards**
    Includes but is not limited to:
    *Exposure to extreme temperatures (heat, cold, or sudden weather changes).*

    *Rugged or uneven terrain, including volcanic, coastal, forested, or mountainous areas.*

    *Slips, trips, and falls while hiking, wading, or working in field sites.*

    *Encounters with wildlife or insects (including stings, bites, or allergic reactions).*

    *Contact with sharp objects, coral, or contaminated water.*

    **Please initial to acknowledge and accept the above risks:**

    ______________________________

11. **Travel & Transportation Risks**
    Includes but is not limited to:

    *Vehicle or boat accidents during transport to and from field sites.*

    *SCUBA diving- or snorkeling-related injuries (e.g., barotrauma, decompression sickness).*

    *Marine hazards (e.g., strong currents, waves, jellyfish, or boat strikes).*

    *Risks associated with air travel, including delays, lost baggage, or emergency landings.*

    **Please initial to acknowledge and accept the above risks:**

    ______________________________

12. **Health & Medical Risks**
Includes but is not limited to:

*Limited access to immediate medical care in remote areas.*

*Illness due to poor sanitation, foodborne pathogens, or waterborne diseases.*

*Fatigue, dehydration, or heatstroke from long days in the field.*

*Exacerbation of pre-existing medical conditions.*

**Please initial to acknowledge and accept the above risks:**

______________________________

13. **Lab & Sample Handling Hazards**
Includes but is not limited to:

*Exposure to chemicals or reagents during sample processing, handling and/or experimental workflow.*

*Biological risk from handling environmental samples, including unknown microbes.*

*Accidental cuts or punctures from sharps or glassware.*

**Please initial to acknowledge and accept the above risks:**

______________________________

14. **The Participant voluntarily assumes all such risks, whether known or unknown. Please initial to acknowledge and accept the above:**

______________________________

#### PARTICIPATION WAIVER AND LEGAL TERMS

*The following section outlines important legal and safety terms related to your participation in The Two Frontiers Project's research activities. These terms are designed to clarify expectations, acknowledge potential risks, and ensure that all participants understand their rights and responsibilities. Please read each statement carefully and provide your initials to confirm your understanding and agreement. If you have any questions, please*

15. **Liability**

    To the fullest extent permitted by law, the Participant hereby releases, waives, and discharges 2FP, its directors, officers, employees, affiliates, partners, sponsors, funders, and agents (collectively, the "Released Parties") from any and all claims, liabilities, demands, actions, or causes of action arising from or related to their involvement with 2FP, including but not limited to claims related to personal injury, property damage, financial loss, or any other harm.

    **Please initial to acknowledge and accept the above:**

    ______________________________

16. **Medical Fitness**

    I certify that I am physically and mentally fit to participate in this expedition. If I have a pre-existing medical condition, I understand that I must obtain written approval from a licensed medical professional before participating. I have disclosed any relevant medical conditions to the organizers and understand that I am responsible for managing my health during the expedition.

    **Please initial to acknowledge and accept the above:**

    ______________________________

17. **Indemnification**

The Participant agrees to indemnify, defend, and hold harmless the Released Parties from and against any claims, damages, losses, liabilities, and expenses (including reasonable attorneys' fees) arising out of or related to the Participant's involvement with 2FP.

**Please initial to acknowledge and accept the above:**

______________________________

18. **Consent to Medical Treatment**

I authorize The Two Frontiers Project to provide or arrange for medical treatment deemed necessary during the expedition. I agree to be financially responsible for any medical services provided.

**Please initial to acknowledge and accept the above:**

______________________________

19. **No Guarantee of Outcome**

The Participant acknowledges that 2FP makes no representations or guarantees regarding the outcomes, findings, or benefits of participation. Any research, data, or results derived from 2FP activities are subject to scientific interpretation and external factors beyond 2FP's control.

**Please initial to acknowledge and accept the above:**

______________________________

20. **No Employment of Compensation Relationship**

The Participant understands and agrees that participation in 2FP activities does not create an employment, partnership, or compensation-based relationship. Unless otherwise agreed upon in writing, all contributions to 2FP are voluntary, and no financial compensation will be provided.
**Please initial to acknowledge and accept the above:**

______________________________

21. **Compliance with Laws and Safety Protocols**

The Participant agrees to comply with all applicable laws, safety guidelines, institutional policies, and ethical standards while participating in 2FP activities. Failure to do so may result in termination of involvement with 2FP.
**Please initial to acknowledge and accept the above:**

______________________________

22. **Consent for Data and Sample Collection**

The Participant acknowledges that 2FP may collect environmental and biological samples for research purposes. Any samples or data collected during the expedition become the property of 2FP and may be used for scientific study, publication, or educational purposes. The Participant understands they will not receive compensation for any data or sample contributions.

**Please initial to acknowledge and accept the above:**

______________________________

23. **Governing Law**

This Agreement shall be governed by and construed in accordance with the laws of the countries and/or regions of travel, without regard to its conflict of law provisions.
**Please initial to acknowledge and accept the above:**

_______________________________

24. **Severability**

If any provision of this Agreement is found to be invalid or unenforceable, the remaining provisions shall remain in full force and effect.
**Please initial to acknowledge and accept the above:**

_______________________________

25. **Acknowledgement and Understanding**

I have read this waiver and fully understand its terms. I acknowledge that I am signing it freely and voluntarily and intend my signature to be a complete and unconditional release of all liability.

**Please initial to acknowledge and accept the above:**

_______________________________

This content is neither created nor endorsed by Google.

Google Forms
