## Supplementary File S3 Example expedition planning and execution package (completed template) V11 for "The Two Frontiers Project Field Handbook and OpenTools: Standardizing microbial fieldwork for biobank-scale sequencing and culturomics": README_08.docx

```
Folder: 08_ADDITIONAL_DATA
 Project: The Two Frontiers Project (2FP)
 Expedition: [Expedition Name]
This folder contains supplementary materials that do not fit cleanly into other standardized folders.
Typical contents include:
```

- Insurance documentation
- Shipping or dry ice documentation
- Special case records
- Miscellaneous supporting files required for the expedition

Use this folder sparingly to avoid fragmentation of core operational data.
