## Supplementary File S3 Example expedition planning and execution package (completed template) V11 for "The Two Frontiers Project Field Handbook and OpenTools: Standardizing microbial fieldwork for biobank-scale sequencing and culturomics": Supplementary File S3_ Example expedition planning and execution package (completed template).docx

*Supplementary File S3: Example expedition planning and execution package (completed template), including team logistics, sampling metadata, inventory tracking, and post-expedition debrief*

This provides a completed example of the Two Frontiers Project (2FP) expedition planning and execution framework. It is intended to demonstrate how the protocols, templates, and workflows described in the 2FP Field Handbook are operationalized during a real-world field campaign.

The materials included here represent a worked example, rather than prescriptive requirements, and are designed to be adaptable to a wide range of expedition types, environments, and team sizes.

This supplement consists of the following components:

- S3a — Team and Travel (Excel)
   Pre-expedition planning of personnel, roles, travel logistics, accommodations, and emergency contacts.
- S3b — Sampling and Metadata (Excel)
   Standardized collection of sampling information, site identifiers, environmental metadata, and sample accessioning fields aligned with the 2FP Living Database.
- S3c — Inventory and Transport (Excel)
   Tracking of field equipment, consumables, sample containers, cold-chain requirements, and transport logistics across the expedition lifecycle.
  S3d — Budget and Expenses (Excel)
   High-level budget planning and expense tracking for expedition costs. Values shown are illustrative and may be generalized.
- S3e — Concept of Operations (Document)
   A day-by-day run-of-show outlining expedition objectives, timelines, locations, personnel responsibilities, and communication plans.
- S3f — Debrief (Excel)
   Post-expedition documentation capturing outcomes, deviations from plan, challenges encountered, and recommendations for future field campaigns.

Additionally, the following folders contain organized and relevant information for expedition planning:

- 01_BACKGROUND_RESEARCH
  - The 01_Background_Research folder contains preparatory materials assembled during the early planning stages of an expedition. While this folder is typically empty in the base 2FP expedition template, it serves as a flexible holding space for contextual information that informs site selection, scientific framing, partner outreach, and proposal development.
  - In this example expedition, the folder includes:
- Literature and reference materials relevant to the target environment and research questions
- Planning documents outlining candidate dates, sites, and logistical constraints
- Proposal and briefing documents prepared for funders, collaborators, or internal review
- Presentation materials (e.g., slides) developed for planning meetings or partner discussions
- 02_FIELD (MAPS, GIS)
  - The 02_FIELD (MAPS, GIS) folder contains spatial materials used to support site selection, navigation, and field execution. These files provide geographic context and practical reference for sampling locations before and during the expedition.
  - In this example expedition, the folder includes:
    - Regional and site-level maps provided by local partners or public sources
    - GIS files and coordinate lists used for locating sampling sites
    - Route planning maps and access paths
    - Annotated screenshots or exported visuals shared during planning
- 03_PERMITTING
  - The 03_PERMITTING folder contains regulatory, legal, and access documentation required to conduct fieldwork at the selected sites. This folder supports compliance with institutional, local, and national requirements.
  - In this example expedition, the folder includes:
    - Permits and access agreements
    - Memoranda of cooperation or letters of authorization
    - Institutional forms related to equipment or field activities
    - Email correspondence and draft documents associated with the permitting process
- 04_PROTOCOLS_EXPERIMENTS
  - The 04_PROTOCOLS_EXPERIMENTS folder contains scientific protocols and reference materials used during sampling and experimental activities. These materials support standardized and reproducible field methods.
  - In this example expedition, the folder includes:
    - Sampling and processing protocols
    - Instrument or sensor documentation
    - Relevant excerpts of the 2FP Field Handbook
    - Reference figures or diagrams used during fieldwork
- 05_PHOTOS
  - The 05_PHOTOS folder contains photographic documentation collected throughout the expedition. These images support scientific records, field notes, and communication or outreach activities.
  - In this example expedition, the folder includes:
    - Site photos documenting sampling locations
    - Sample and workflow photo
    - Lab notebook or procedural images
    - Drone imagery and press or outreach photos
- 06_BILLING_RECEIPTS
  - The 06_BILLING_RECEIPTS folder contains financial documentation associated with expedition-related expenses. These files support reimbursement, reporting, and internal accounting.
  - In this example expedition, the folder includes:
    - Travel receipts
    - Lodging invoices
    - Equipment and supply receipts
    - Other reimbursable expense records
- 07_FORMS
  - The 07_FORMS folder contains required forms related to expedition participation and compliance. These documents support safety, liability, and institutional requirements.
  - In this example expedition, the folder includes:
    - Field participation waivers
    - Safety or activity-specific forms
    - Diving or training-related documentation
- 08_ADDITIONAL_DATA
  - The 08_ADDITIONAL_DATA folder contains supplementary materials that do not fit cleanly within other standardized folders but are necessary for expedition support.
  - In this example expedition, the folder includes:
    - Insurance documentation
    - Shipping or dry ice records
    - Other supporting materials required for the expedition

Together, these files span the full expedition lifecycle:

1. Pre-expedition planning (team, travel, permitting, inventory, and budget)
2. Field execution (daily operations, sampling, metadata capture)
   Post-expedition review (debrief, lessons learned, and data reconciliation)

While each file can be used independently, they are designed to function as an integrated system supporting reproducibility, operational clarity, and standardized data collection.

This example has been completed using a real or representative expedition scenario. Personally identifiable information and sensitive logistics have been removed or generalized. The structure, fields, and workflows reflect best practices developed by 2FP across multiple field campaigns.

All templates used to generate this example are maintained as open-source, version-controlled resources in the Two Frontiers Project Field Handbook repository:
https://github.com/two-frontiers-project/2FP-Field-Handbook

Users are encouraged to adapt these materials to their specific research, regulatory, and logistical needs.
