## Supplementary File S3 Example expedition planning and execution package (completed template) V11 for "The Two Frontiers Project Field Handbook and OpenTools: Standardizing microbial fieldwork for biobank-scale sequencing and culturomics": Supplementary_File_S3e_Concept_of_Operations.doc

Expedition Communication – Concept of Operations

Topic: CARBON9 – Spence Hot Springs, New Mexico

Description:
 Run of show and operational overview for a pilot terrestrial geothermal sampling expedition conducted as part of the 2FP Carbon program.

Timeline:
 May 16–18, 2026

Purpose of Trip:
 In May 2026, The Two Frontiers Project (2FP) will conduct a pilot terrestrial field expedition to Spence Hot Springs in the Santa Fe National Forest, New Mexico. The goal of the expedition is to collect microbial mat, sediment, and water samples from a low-temperature geothermal environment, record standardized environmental metadata, and generate baseline data to inform future geothermal sampling efforts within the Valles Caldera region.

Primary Research Location:
 Spence Hot Springs
 Santa Fe National Forest / Valles Caldera region
 New Mexico, USA

Expedition Summary:

Friday, May 16, 2026
 Team arrival in Santa Fe, New Mexico.
 Vehicle pickup, procurement of final field supplies, and overnight lodging near the field site.
 Evening planning meeting to review sampling locations, safety considerations, and field workflow.

Saturday, May 17, 2026
 Primary field sampling day.
 Team accesses Spence Hot Springs via established hiking trails.
 Sampling of microbial mats, shallow sediments, and spring water conducted using hand tools only.
 Environmental metadata recorded at each sampling location.
 Samples stored in insulated containers and transported back to lodging for temporary storage.

Sunday, May 18, 2026
 Optional backup sampling window or site revisits if needed.
 Final sample checks, packing, and preparation for transport to laboratory facilities.
 Team departure from Santa Fe.
 Conclusion of expedition.

Sampling Day Overview (May 17, 2026)

Morning
 Depart lodging for field site.
 Conduct site safety assessment and confirm sampling locations.
 Set up sampling equipment and metadata recording materials.

Midday
 Collect microbial mat, sediment, and water samples.
 Record GPS coordinates, water temperature, pH, visual site observations, and weather conditions.
 Photograph sampling locations and workflows for documentation.

Afternoon
 Pack and secure samples for transport.
 Restore sampling areas as appropriate.
 Return to lodging.

Evening
 End-of-day debrief.
 Review sampling outcomes and note any deviations or observations relevant to future campaigns.
